# Molecular docking between human TMPRSS2 and the serine protease Kunitz-type inhibitor rBmTI-A

**DOI:** 10.1101/2022.03.13.484191

**Authors:** Lívia de Moraes Bomediano Camillo, Sergio Daishi Sasaki

## Abstract

SARS-CoV-2 entrance into host cells is dependent of ACE2 receptor and viral protein S initiation by serine protease TMPRSS2. Cleavage of coronavirus protein S at the junctions Arg685/Ser686 and Arg815/Ser816 leads to the production of the S1/S2 and S2’ fragments needed for the fusion of viral and cell membranes. Studying and identifying serine protease inhibitors is an important step towards the development of candidate drugs to prevent SARS-CoV-2 infection. It has already been stablished that camostat mesylate, a serine protease inhibitor, is capable of blocking TMPRSS2 activity and prevent SARS-CoV-2 entrance into host cells. In this work, the interaction between the two domains of Kunitz-type serine protease inhibitor rBmTI-A and TMPRSS2 was studied through molecular docking. rBmTI-A domain 2 (P1 site Leu84) had the best complex results with predicted binding affinity of -12 Kcal.mol^-1^ and predicted dissociation constant at 25°C of 1.6 nM. The results suggest that rBmTI-A is capable of binding TMPRSS2 cleavage site at the junction Arg815/Ser816 using essentially the same residues that camostat mesylate.

## Background

Human transmembrane protease serine-2 (TMPRSS2) is a serine protease highly expressed in prostate cancer [1–4] and its role has been described in other tumorigenesis processes and cancer pain [5–7]. TMPRSS2 also plays an important role in the infection process caused by severe acute respiratory syndrome coronavirus (SARS-CoV) [8,9] and severe acute respiratory syndrome coronavirus 2 (SARS-CoV-2) [10–13].

The recognition of receptors by viruses is a key step in viral infectivity and pathogenesis. Therefore, it is a target for the development of drugs and vaccines. In previous studies with other types of coronaviruses such as SARS-CoV, the interaction between viral proteins and cell receptors has been established [14–17]. The viral surface protein S (spike) mediates the entry of the virus into the host cell through its interaction with the angiotensin-converting enzyme 2 (ACE2, angiotensin-converting enzyme 2). Protein S has a receptor-binding domain (RBD) that specifically recognizes ACE2 as its receptor [18,19].

SARS-CoV-2 entry into host cells also depends on the ACE2 receptor and viral protein S initiation by serine protease TMPRSS2 [10,11,13]. Other studies have demonstrated and characterized the interaction of SARS-CoV-2 with the ACE2 receptor [20,21]. Furthermore, Hoffmann et al., 2021 showed that the camostat mesylate drug, a serine protease inhibitor used to treat chronic pancreatitis [23] was able to block the action of TMPRSS2 and, therefore, block the entry of the virus into host cells. Baughn et al., 2020 and Wettstein et al., 2021 demonstrated that serine protease inhibitor alpha-1 antitrypsin is also capable to inhibit the action of TMPRSS2 and stop the development of the SARS-CoV-2 infection. These results reflect the importance of studying and identifying more serine protease inhibitors as potential drug candidates to prevent SARS-CoV-2 infection [10,24].

rBmTI-A is a recombinant Kunitz-BPTI serine protease inhibitor with inhibitory action against trypsin, human plasma kallikrein (HuPK), HNE and human plasmin [25,26]. The inhibitor was initially obtained from the extract of larvae of the bovine tick *Rhipicephalus microplus* [25–27] and it has two Kunitz-BPTI domains, with a total molecular weight of 13.87 kDa. Domain 1 has the amino acid Arg at its P1 site with inhibitory action against trypsin and HuPK. Domain 2 has the amino acid Leu in its reactive site with inhibitory action against HNE [26].

The serine protease inhibitor rBmTI-A has shown important experimental results in the context of pulmonary emphysema and lung tissue inflammation [27–29]. Given the health emergency imposed by SARS-CoV-2, responsible for a disease that mainly affects the lung tissue and the relationship between the interaction of its protein S with the ACE2 receptor through an initialization of protein S by the serine protease TMPRSS2, the study of the interaction of TMPRSS2 and serine protease inhibitor rBmTI-A can lead to development of a therapeutic potential drug against COVID-19 disease.

## Methods

### Structure data mining

TMPRSS2 structure was previous modeled by Hussein et al., 2020 [30] and made publicly available by the authors. rBmTI-A structure was modeled by our group in previous work [31]. Molecular dynamic simulations were performed to flexibilize both structures in water as solvent for 100 nanoseconds using NAMD v.2.14 [32] under default parameters.

### Molecular docking

The prediction of the interaction interface between TMPRSS2 and rBmTI-A was made using HADDOCK v.2.2 webserver [33,34]. TMPRSS2 catalytic triad (His296, Asp345 and Ser441) and amino acids already described as involved in the substrate binding sites (Asp435, Ser460 and Gly462) [30] were defined as active residues. rBmTI-A domain 1 P1 site Arg24 and domain 2 P1 site Leu84 were defined as active residues. The passive residues were defined as the surrounded amino acids in both cases. The cluster with the best HADDOCK score was chosen for further pose analysis. PRODIGY webserver v.1.0 [35] and Ligplot^+^ v.2.2 [36] were used to assess binding affinity energy and different types of interactions. Molecular dynamic simulations were performed to flexibilize final complex structures in water as solvent for 100 nanoseconds using NAMD v.2.14 [32] under default parameters. All structures were visualized using Pymol v.2.4.1 (The PyMOL Molecular Graphics System, Version 2.0 Schrödinger, LLC) and VMD v.1.9.4a51 [37].

## Results

### Interaction of TMPRSS2 and serine protease inhibitor rBmTI-A

The interaction complexes between TMPRSS2 and P1 sites 1 (Arg24) and 2 (Leu84) are shown in Figure 1 and Figure 2. Amino acids participating in the intermolecular interaction are shown as orange sticks in TMPRSS2 and as cyan sticks in rBmTI-A. The binding affinity of the complex model with the interaction occurring at rBmTI-A P1 site 1 (Arg24) is -10.4 kcal mol^-1^ with Kd at 25°C of 0.25 nM and at 37°C of 0.45 nM. The binding affinity of the complex model with the interaction occurring at rBmTI-A P1 site 2 (Leu84) is -12 kcal mol^-1^ with Kd at 25°C of 1.6 nM and at 37°C of 3.5 nM.

**Figure 1.**
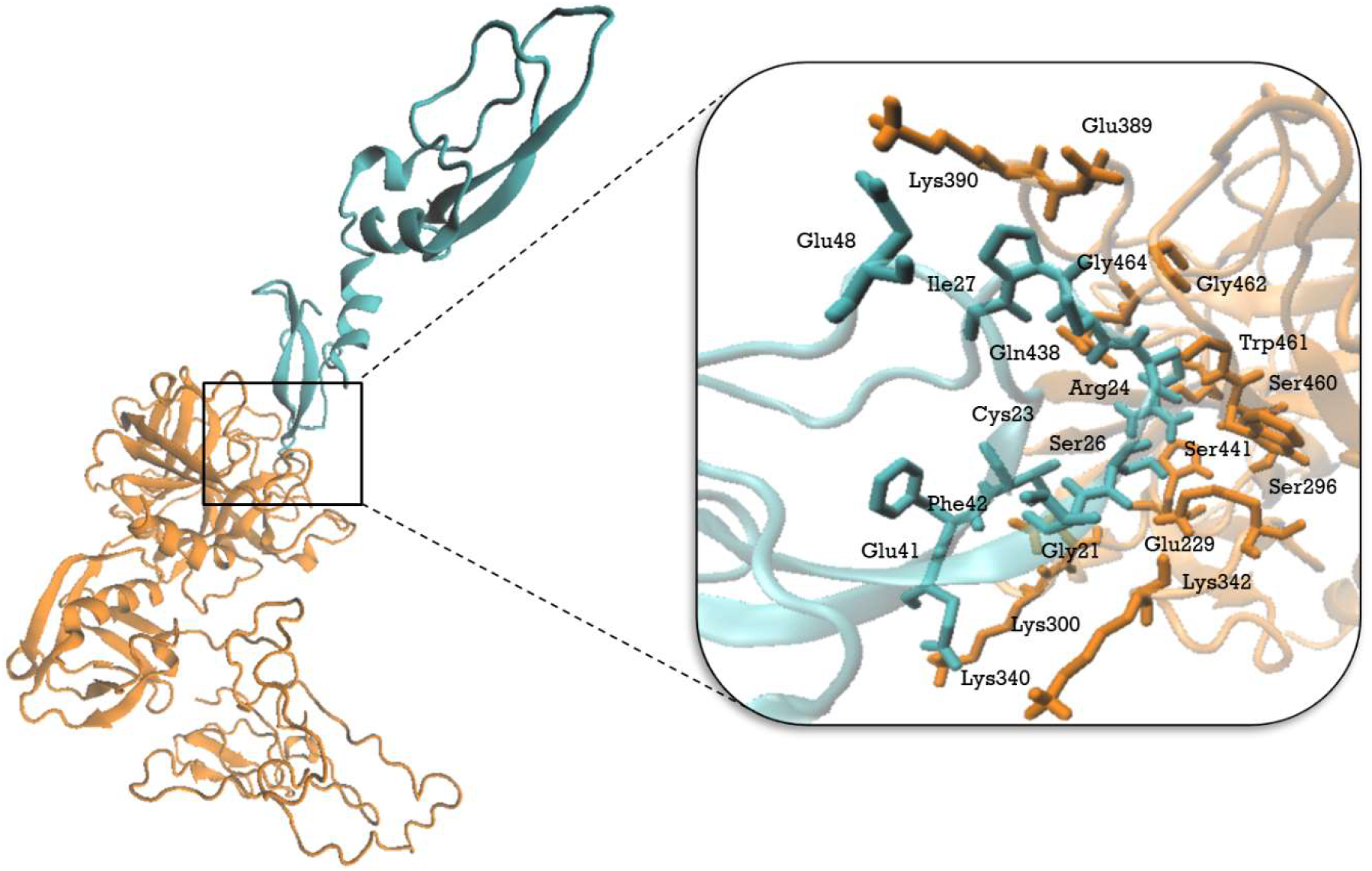
TMPRSS2 (orange) and rBmTI-A (cyan) complex model with interaction in rBmTI-A P1 site Arg24. Amino acids participating in the intermolecular interaction (TMPRSS2 residues as orange sticks and rBmTI-A residues as cyan sticks) are specified in the box. Binding affinity of the complex model is -10.4 kcal mol^-1^ with Kd at 25°C = 0.25 nM and at 37°C = 0.45 nM.

**Figure 2.**
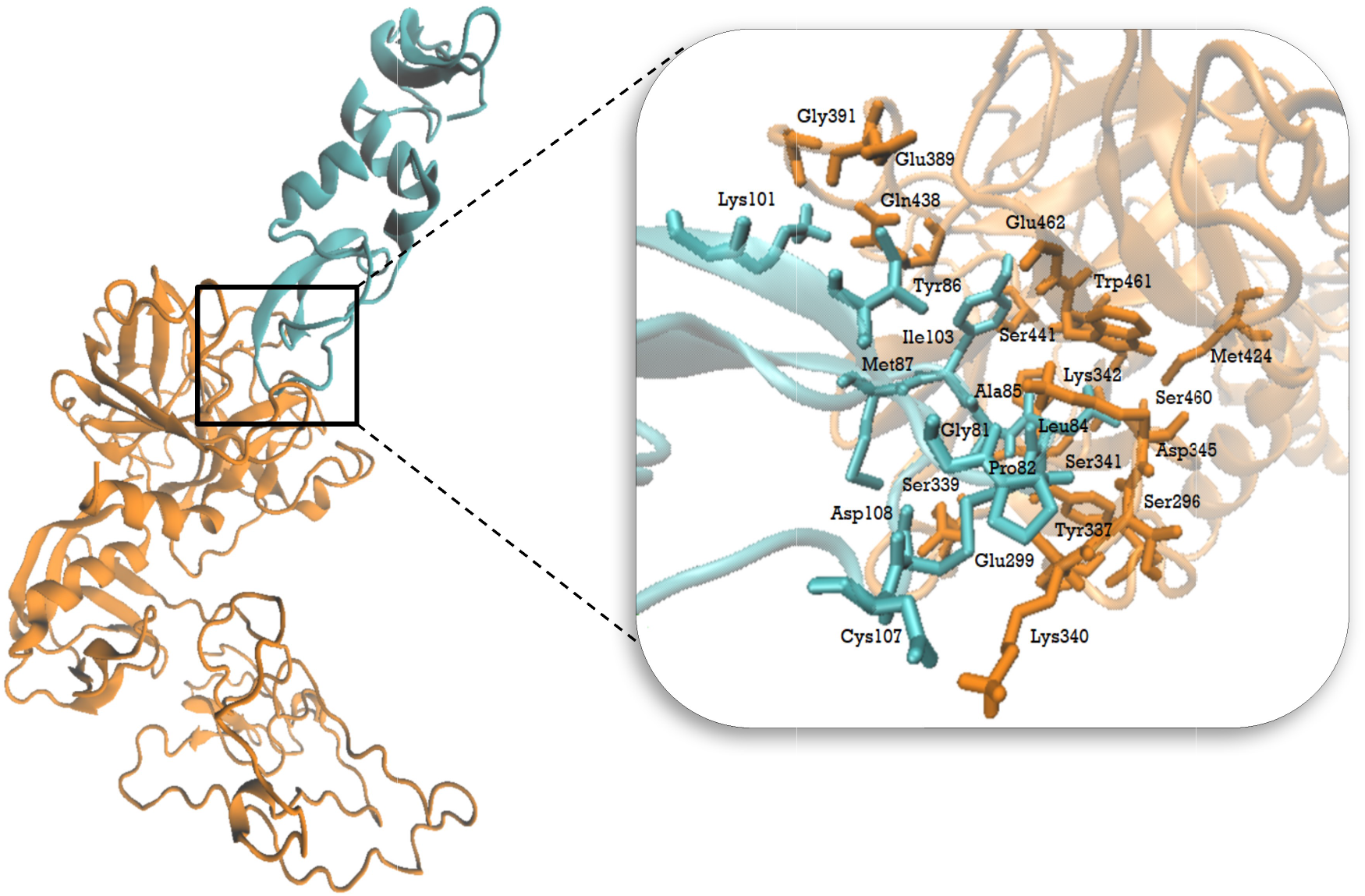
TMPRSS2 (orange) and rBmTI-A (cyan) complex model with interaction in rBmTI-A P1 site Leu84. Amino acids participating in the intermolecular interaction (TMPRSS2 residues as orange sticks and rBmTI-A residues as cyan sticks) are specified in the box. Binding affinity of the complex model is -12 kcal mol^-1^ and Kd at 25°C = 1.6 nM and at 37°C = 3.5 nM.

TMPRSS2 interface residues involved in proteolytic cleavage of the Sars-CoV-2 spike protein have been determined and the proteolytic cleavage sites has been mapped at the junction of Arg685/Ser686 and Arg815/Ser816 [30]. Table 1 shows the comparison between TMPRSS2 interface residues found in the interaction with SARS-CoV-2 cleavage sites 1 and 2 [30] with camostat mesylate inhibitor [38] and rBmTI-A inhibitor P1 sites 1 and 2.

**Table 1.** TMPRSS2 interface residues in the interaction with SARS-CoV-2 cleavage sites 1 and 2, with camostat mesylate inhibitor and rBmTI-A inhibitor P1 sites 1 and 2.

| **TMPRSS2 interface**<br>**residues with SARS-**<br>**CoV-2**<br>**cleavage site 1\***<br>**(Arg685/Ser686)** | **TMPRSS2 interface**<br>**residues with SARS-**<br>**CoV-2**<br>**cleavage site 2\***<br>**(Arg815/Ser816)** | **TMPRSS2**<br>**interface**<br>**residues**<br>**with**<br>**camostat**<br>**mesylate\*\*** | **TMPRSS2**<br>**interface**<br>**residues**<br>**with rBmTI-**<br>**A**<br>**(Site Arg24)** | **TMPRSS2**<br>**interface**<br>**residues**<br>**with rBmTI-**<br>**A**<br>**(Site Leu84)** |
| --- | --- | --- | --- | --- |
| Val275 | Val278 | Val280 | His296 | His296 |
| Gln276 | Val280 | Ala295 | Glu299 | Glu299 |
| His296 | His296 | His296 | Lys300 | Ser339 |
| Glu299 | Glu299 | Glu299 | Lys340 | Tyr337 |
| Lys300 | Lys300 | Tyr337 | Lys342 | Lys340 |
|  |  | Lys342 |  |  |
| Thr387 | Asp338 | Asp345 | Glu389 | Lys342 |
| Glu388 | Lys340 | Gly391 | Lys390 | Thr341 |
| Lys390 | Thr341 | Lys392 | Gln438 | Asp345 |
| Gly391 | Lys342 | Thr393 | Ser441 | Glu389 |
| Arg413 | Glu389 | Gln438 | Ser460 | Gly391 |
| Tyr414 | Gly391 | Asp440 | Trp461 | Gln438 |
| Gln431 | Lys392 | Ser441 | Gly462 | Ser441 |
| Asn433 | Leu419 |  | Gly464 | Ser460 |
| Ala466 | Gln438 |  |  | Gly462 |
| Tyr469 | Ser441 |  |  | Trp461 |
|  | Ser460 |  |  |  |
|  | Gly462 |  |  |  |
|  | Cys465 |  |  |  |
|  | Lys467 |  |  |  |

\*[30,39], \*\*[38].

Tables 2 and 3 show the main type of interactions between rBmTI-A P1 sites and TMPRSS2 residues. Figure 3 shows the comparison between rBmTI-A and TMPRSS2 interfacial contacts at rBmTI-A P1 sites. At the interaction in rBmTI-A P1 site 1 (Arg24), Arg24 forms hydrogen bonds with TMPRSS2 residues Gln438, Ser460 and Gly462 and hydrophobic interaction with His296 and Ser441. Whereas at the interaction in rBmTI-A P1 site 2 (Leu84), Leu84 forms a hydrogen bond with TMPRSS2 residue Tyr337 and hydrophobic interaction with Tyr337, Met424 and Trp461. These results indicate that rBmTI-A is capable of binding to the residues of TMPRSS2 interacting at the interface with both SARS-CoV-2 spike protein cleavage sites and with camostat mesylate. Furthermore, these findings shows that rBmTI-A P1 site Leu84 has more interfacial contacts with TMPRSS2, with better energy binding affinity and dissociation constant.

**Table 2.** Corresponding residues and main interactions type between rBmTI-A P1 site Arg24 and TMPRSS2.

| Interaction type | TMPRSS2 | rBmTI-A<br>(Arg24) |
| --- | --- | --- |
| **Intermolecular** | Gly462 | Arg24 |
| **Hydrogen Bonds** | Lys342 | Ser26 |
|  | Gln438 | Gly21 |
|  | Gln438 | Cys23 |
|  | Ser460 | Arg24 |
|  | Glu299 | Thr43 |
|  | Lys300 | Glu41 |
|  | Lys390 | Glu48 |
|  | Gln438 | Arg24 |
| **Intermolecular** | Trp461 | Ala25 |
| **Hydrophobic** | His296 | Arg24 |
| **Interactions** | Ser441 | Arg24 |

**Table 3.**
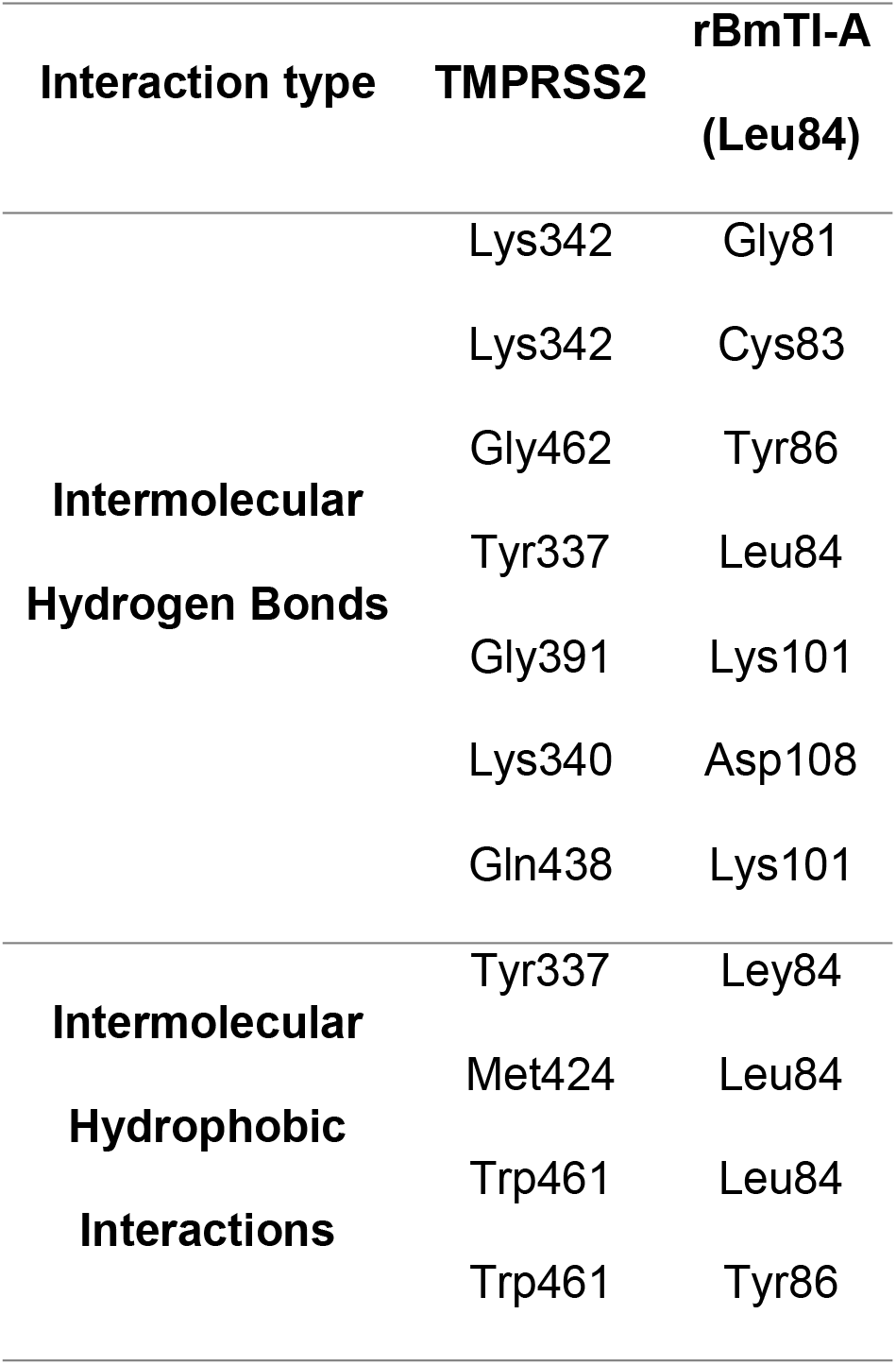
Corresponding residues and main interactions type between rBmTI-A P1 site Leu84 and TMPRSS2.

**Figure 3.**
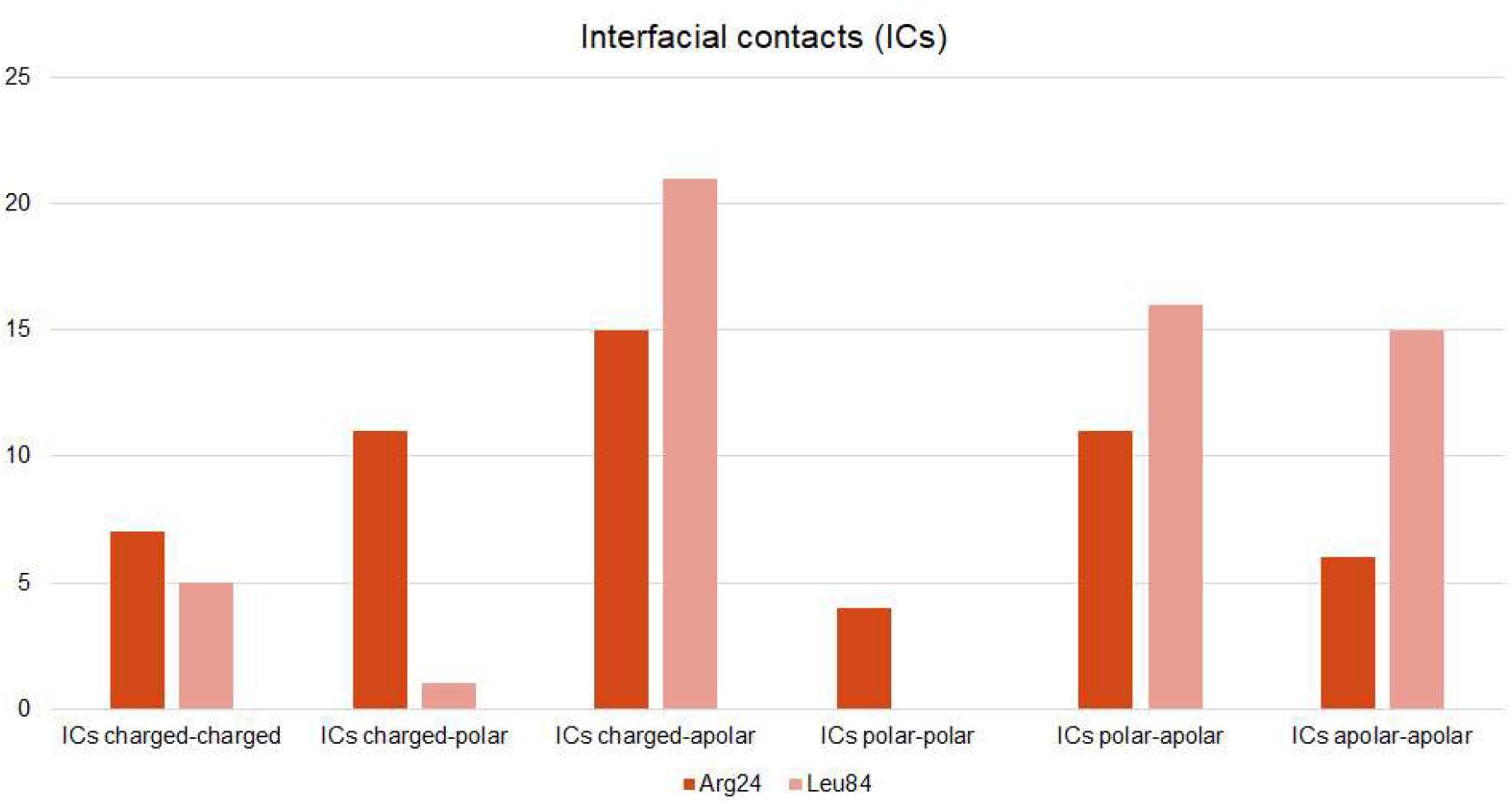
Interfacial contacts (ICS) between rBmTI-A sites (Arg24 in red, Leu84 in salmon) and TMPRSS2. Interaction at P1 site Arg24 has a total of 54 interfacial contacts with TMPRSS2, while at P1 site Leu84 has 58.

## Discussion

In late February 2003, the first outbreak of SARS-CoV viral respiratory disease emerged in China and spread to other countries (World Health Organization. Retrieved from *www.who.int/health-topics/severe-acute-respiratory-syndrome*). Following the emergence of the outbreak, Li et al identified the functional receptor for SARS-CoV as the angiotensin-converting enzyme 2 (ACE2) [18]. ACE2 is a type I membrane protein mainly expressed in lungs, heart, kidneys and intestine) [40,41]. ACE2 primary physiological role is the maturation of angiotensin, a peptide hormone that controls vasoconstriction and blood pressure [40]. Li et al and Wu et al demonstrated how ACE2 is capable of efficiently bind the S1 domain of the SARS-CoV spike protein (S) and enable host and viral membranes fusion [16,18].

The entry of SARS-CoV into the host cell also depends on the initialization of viral protein S by cellular proteases which implies cleavage of protein S at the junctions Arg685/Ser686 and Arg815/Ser816, which leads to the production of the S1/S2 and S2’ fragments needed for the fusion of viral and cell membranes [14–17,30]. The initialization of viral protein S is done by TMPRSS2 [8,9,42,43]. The TMPRSS2 serine protease is a type II transmembrane serine protease that are expressed in many human tissues, such as prostate, stomach, salivary gland, mammary gland, pancreas, kidney, liver, small intestine, lung, trachea, fetal kidney, fetal lung, eye and airway epithelial cells [44,45]. SARS-CoV-2, responsible for the ongoing disease outbreak, is closely related to SARS-CoV and both share the same mechanisms of host cell fusion and entrance [19,30,46].

The coronavirus spike protein is a 1273 amino acid long homo-trimeric glycoprotein that has two major subunits: S1, involved in receptor recognition and binding and membrane anchored S2, involved in the mediation of the fusion of the viral and cell host membranes [47–49]. S protein is first cleaved at S1/S2 site, assuming a prefusion conformation. Then, a second cleavage occurs at S2’ site which leads to an irreversible conformational change that result in membrane fusion and viral entrance into the host cell [46,50–54]. Receptor binding and membrane fusion are the initial critical step in the coronavirus infection cycle, and it is as primary target for therapeutical approaches.

Several studies are advancing around the activity of serine protease inhibitors toward TMPRRS2. Camostat mesylate and nafamostat mesylate are two serine protease inhibitors that have shown activity blocking TMPRSS2 action in SARS-CoV and MERS-CoV infections [55,56]. Camostat mesylate is a serine protease inhibitor developed for the treatment of chronic pancreatitis and postoperative reflux esophagitis, but other usage has been reported in cancer treatment and fibrosis inhibition [57–59]. Sivaraman et al and Sonawane et al have characterized the interaction between serine protease inhibitor camostat mesylate and TMPRRSS2 [38,39]. The binding energy of this interaction is - 6,23 kcal.mol^-1^ and its inhibition constant (Ki) is 26.98 nM.

Camostat and TMPRSS2 residues form five hydrogen bonds at the interaction interface and TMPRSS2 amino acid residues Ile381, His296 and His279 are involved in hydrophobic interactions. Asn398, Gly282, His296 and Cys281 provide Van der Waals interactions [39]. Hoffmann et al confirmed that camostat also has inhibitory effects on the SARS-CoV-2 in TMPRSS2-expressing human cells [22]. Yamamoto et al also confirmed that nafamostat can block SARS-CoV-2 fusion at a lower concentration compared to camostat [60]. These results indicated that studies of serine protease inhibitors as potential drug candidates are crucial in the development of a treatment for COVID-19.

Serine protease inhibitors of the Kunitz-BPTI family have one or more Kunitz-BPTI domains being characterized by a conserved spacing between cysteine residues, a typical pattern of disulfide bond [61,62]. The Kunitz domain has a secondary alpha + beta-type structural fold, where alpha helices and beta sheets occur separately along the skeleton. The domain is characterized by the presence of three highly conserved disulfide bonds that are necessary for stabilization of these inhibitors native conformation [61,62]. The bovine pancreatic trypsin inhibitor (Bovine Pancreatic Trypsin Inhibitor - BPTI) is the typical model of Kunitz-BPTI type inhibitors. BPTI has broad specificity, being able to inhibit trypsin, chymotrypsin and elastase-type serine proteases [63].

Tanaka et al characterized a Kunitz-type inhibitor from tick larvae [64]. BmTI-A is an 18 kDa inhibitor with activity on several serine proteases, such as trypsin, HNE, human plasma kallikrein, human plasmin and chymotrypsin [25,26,64]. The recombinant BmTI-A (rBmTI-A) is a serine protease inhibitor of the Kunitz-BPTI type with inhibitory action against trypsin, human plasma kallikrein, HNE and human plasmin in nM range [26]. Like the BmTI-A inhibitor, it has two Kunitz-BPTI domains and a total molecular weight of 13.87 kDa. Domain 1 has the amino acid Arg at its P1 site with an inhibitory action against trypsin and HuPK and domain 2 has the amino acid Leu at its P1 site with an inhibitory action against HNE [26].

rBmTI-A domains have 51-64 residues that form a central anti-parallel beta sheet and a short C-terminal helix. Each domain has six cysteine residues that form three disulfide bonds, resulting in a double loop structure. The active inhibitory binding loop is located between the N-terminal region and the first beta sheet [62]. The structure and thermostability of the rBmTI-A inhibitor were demonstrated by Bomediano Camillo et al [31]. rBmTI-A has been studied by different groups with positive results regarding its effects on chronic obstructive pulmonary disease (COPD), pulmonary emphysema development, chronic allergic pulmonary inflammation and vessel formation [27–29].

The results presented in this work showed that domain 2 of rBmTI-A interacts better with TMPRSS2 cleavage site at the junction Arg815/Ser816 with binding affinity energy of -12 kcal.mol^-1^ and predicted dissociation constant (Kd) in nanomolar scale. rBmTI-A domain 2 and TMPRSS2 residues form seven intermolecular hydrogen bonds and four main hydrophobic interactions. TMPRSS2 residues Lys342, Gly462, Tyr337, Gly391, Lys340 and Gln348 are involved in the hydrogen bonds while residues Tyr337, Met424, Trp461 are involved in the main hydrophobic interactions. The comparison of the amino acids participating in the interface interaction between rBmTI-A and TMPRSS2 versus camostat mesylate and TMPRSS2 are shown in table 1 and these findings suggest that rBmTI-A can interact and block the TMPRSS2 cleavage site essentially at the same positions that camostat mesylate. In addition, rBmTI-A has better predicted binding affinity energy when in a complex with TMPRSS2 than camostat mesylate.

Studies have shown that TMPRSS2 is mainly expressed in the lung tissue, salivary gland, thyroid, gastrointestinal tract, pancreas, kidney, and liver according to RNA and protein expression data available [65]. Matusiak et al studied the differential expression of TMPRSS2 and ACE2 in publicly available data and investigated the effect of smoking behaviour, infections and pollutants in the lung, nasal tissue, bronchial tissue and cell lines [66,67]. TMPRSS2 expression in airway epithelia is highly upregulated by IL-13, a cytokine involved in allergic airway inflammation response, asthma and COPD development [68,69]. It has been shown that IL-13 is also highly expressed in the plasma of COVID-19 patients that require intensive care [70]. These results indicate that TMPRSS2 is greatly expressed in lung tissue inflammation processes and, therefore, COPD contributes to aggravate COVID-19 severity. Since rBmTI-A treatment has shown positive outcome in COPD and lung tissue inflammation [26–29], the study of the interaction and inhibition of TMPRSS2 by rBmTI-A is an important first step in the development of a drug candidate for COVID-19 treatment.

## Conclusion

COVID-19 have no specific treatment established yet. The severity of the disease specially in groups that present comorbidities like COPD, still present a challenge in medical care. The development of potential drug candidates is very important and targeting proteases involved in SARS-CoV-2 entrance to the host cell can help to prevent infection. In this work, we present an initial investigation of the action of the serine protease inhibitor rBmTI-A towards TMPRRS2 as a first step in the development of a potential drug to COVID-19 prevention. The results suggest that rBmTI-A can interact and block TMPRSS2. Further studies should be carried to continue evaluating the possibilities of using rBmTI-A as a drug candidate to COVID-19.

## Acknowledgements

We are grateful to Federal University of ABC for providing the infrastructural and funding support. Also, we are grateful by financial support from Fundação de Amparo à Pesquisa do Estado de São Paulo (FAPESP) 2018/11874-5 and CAPES, financial code 001.

